# Temperament in early childhood is associated with gut microbiota composition and diversity

**DOI:** 10.1101/2023.12.03.569811

**Authors:** Eriko Ueda, Michiko Matsunaga, Hideaki Fujihara, Takamasa Kajiwara, Aya K. Takeda, Satoshi Watanabe, Keisuke Hagihara, Masako Myowa

## Abstract

Temperament is a key predictor of human mental health, and cognitive and emotional development. While human fear behavior is reportedly associated with gut microbiome in infancy, infant gut microbiota changes dramatically during the first five years, when the diversity and composition of gut microbiome are established. This period is crucial for developing human executive functioning, including emotion regulation. Therefore, this study investigated the relationship between temperament and gut microbiota in 284 preschool children aged 3–4 years. Child temperament was assessed by maternal reports of the Children’s Behavior Questionnaire. Gut microbiota (alpha/beta diversity and genera abundance) was evaluated using 16S rRNA sequencing of stool samples. A high abundance of anti-inflammatory bacteria (e.g., Faecalibacterium) and low abundance of inflammatory bacteria (e.g., Eggerthella, Flavonifractor) were associated with higher positive emotionality and reward-seeking (i.e., Surgency/Extraversion, β =0.15, p = 0.013), and lower negative emotionality and behavioral inhibition (i.e., Negative Affectivity, β = -0.17, p = 0.004). Additionally, gut microbiota diversity was associated with a more active approach and exploration (i.e., Impulsivity, a specific aspect of Surgency/Extraversion, β = 0.16, p = 0.008). This study provides insight into the biological mechanisms of temperament and takes important steps toward identifying predictive markers of psychological/emotional risk.

## 1. Introduction

The global prevalence of psychiatric and neurodevelopmental disorders is close to 15% in children up to adolescence [1]; however, signs of these problems are present during infancy [2]. Prevention starting early in life is an urgent concern, and several large cohort studies targeting early intervention have been conducted (e.g., the Healthy Brain and Child Development study [3], the Halland Health and Growth Study [4]).

The gut microbiota has attracted the attention of researchers as a potential early biomarker not only for physical health, such as allergies, asthma, and diabetes [5], but also for mental function [6,7]. Gut bacteria are mediators that modulate the gut-brain axis through the autonomic nervous system, neurotransmitters, and the immune system and are closely related to the brain and behavior [8]. Notably, animal studies have revealed that the manipulation of gut bacteria can alter brain structures and function [9,10].

The relationship between gut microbiota and brain structure/function has been demonstrated in human infancy, albeit in relatively few studies. For example, gut microbiota diversity at one year of age is associated with the activation of functional brain networks important for emotional processing [11], and *Streptococcus* abundance at one month was associated with amygdala volume [12]. Microbiota is also associated with psychological traits in infancy. Temperament, biologically based individual differences in behavioral reactivity to environmental stimuli and self-regulation [13,14], has been investigated in terms of the developmental relationship with infant gut microbiota.

Temperament consists of three components—*Negative Affectivity*, *Surgency/Extraversion*, and *Effortful Control*—each involving different central nervous systems [15]. *Negative Affectivity* reflects the expression of negative emotions and defensive response to potential threats involving limbic and hypothalamus-pituitary-adrenal axis activity. *Surgency/Extraversion* is associated with individual differences in the expression of positive emotions and approach behavior. It involves the activity of the reward system, including the striatum, nucleus accumbens, and amygdala. *Effortful Control* refers to individual differences in the conscious control of emotion and attention. It involves activity in executive functioning networks such as the anterior cingulate cortex and prefrontal cortex. Furthermore, temperament has also attracted researchers’ attention as a predictor of later personality, behavior, and mental health risk [16]. For instance, *Negative Affectivity* in early childhood predicts future internalizing symptoms such as anxiety and depression [17,18].

Most developmental studies investigating the relationship between temperament and microbiota have primarily focused on infancy, and the results have shown inconsistencies. For instance, a study found that 12-month-old infants with lower levels of *Bacteroides* and higher levels of *Lactobacillus* and *Bifidobacterium* exhibited more fearful behavior toward novel non-social stimuli [12]. However, another study reported that low levels of *Lactobacillus* at 12 months were associated with higher *Negative Affectivity* [19] (for more information, refer to the recent review on the association between gut microbiota and temperament [20]).

These inconsistencies may be attributed to the fact that the gut microbiota undergoes significant changes during infancy. A comprehensive study involving children aged 3–48 months reported that the diversity and composition of the top five phyla in the gut microbiota changed significantly between 3 and 14 months of age [21]. Furthermore, even beyond this period, the diversity and composition of the gut microbiota continue to undergo change. However, the main compositions gradually stabilize and approximate those found in adults around 3–5 years after birth [4,21]. Remarkably, this period of stabilization aligns with the significant development of executive functions, including emotional regulation [22,23]. It indicates that early childhood plays a key role in not only establishing the foundation of the future gut microbiota but also in the development of mental functioning in humans. Nevertheless, our understanding of the relationship between gut microbiota and mental functioning during this period remains limited.

This study investigates the relationship between temperament and composition/diversity of gut microbiota in preschool-aged children. It determines if the previously observed link between gut microbiota and temperament in infancy can also be observed in preschool children. We hypothesize that the association between gut microbiota and temperament during the preschool years will differ from that observed in infancy because of the ongoing changes in the composition and diversity of the microbiota.

## 2. Materials and Methods

### 2.1. Participants and procedure

This study is part of a research project in Japan, titled The Principle of Human Social Brain-Mind Development. The participants were recruited from nursery schools throughout Japan that agreed to participate in the study. Written informed consent was obtained from the mothers of the children. Mothers collected stool samples from children and completed questionnaires at home (e.g., temperament, socioeconomic status).

The study collected data on Japanese children aged 0–4 between January and February 2021. A total of 293 children aged 3–4 completed both the stool collection and temperament questionnaires. We excluded nine children for the following reasons: (1) no answer to all or most of the following questions (socioeconomic status, antibiotic use within the last three months, mother’s psychiatric history); and (2) their mother had a psychological disease under medical treatment or was taking medication related to psychiatric disorders. Finally, 284 children were analyzed (mean age = 46.44 ± 6.27 months, 172 boys and 112 girls; for detailed characteristics of participants, see Table 1). This study was approved by the Medical Ethics Committee of Kyoto University (no. R2624) and registered in the UMIN system (UMIN000043945).

**Table 1.**
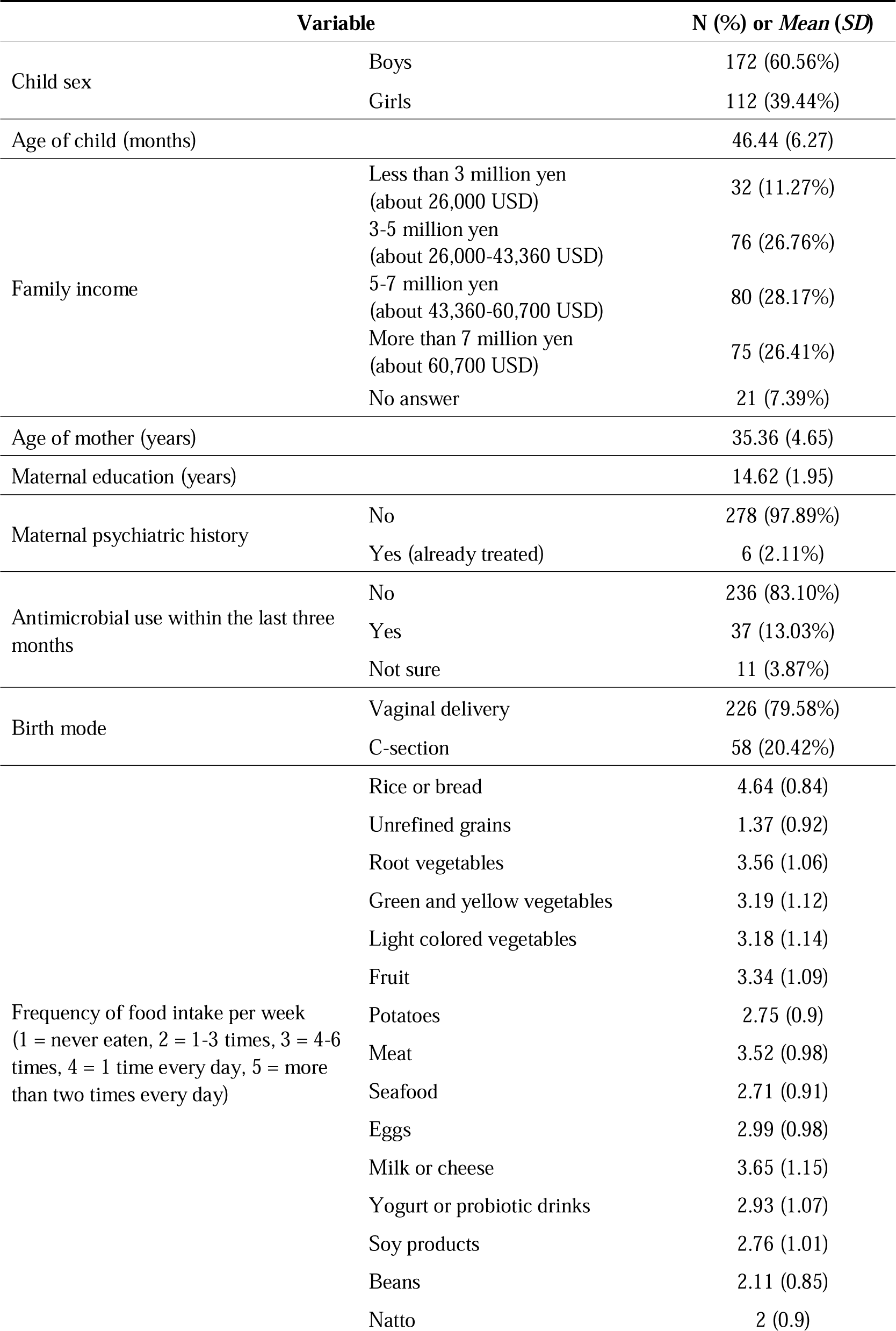
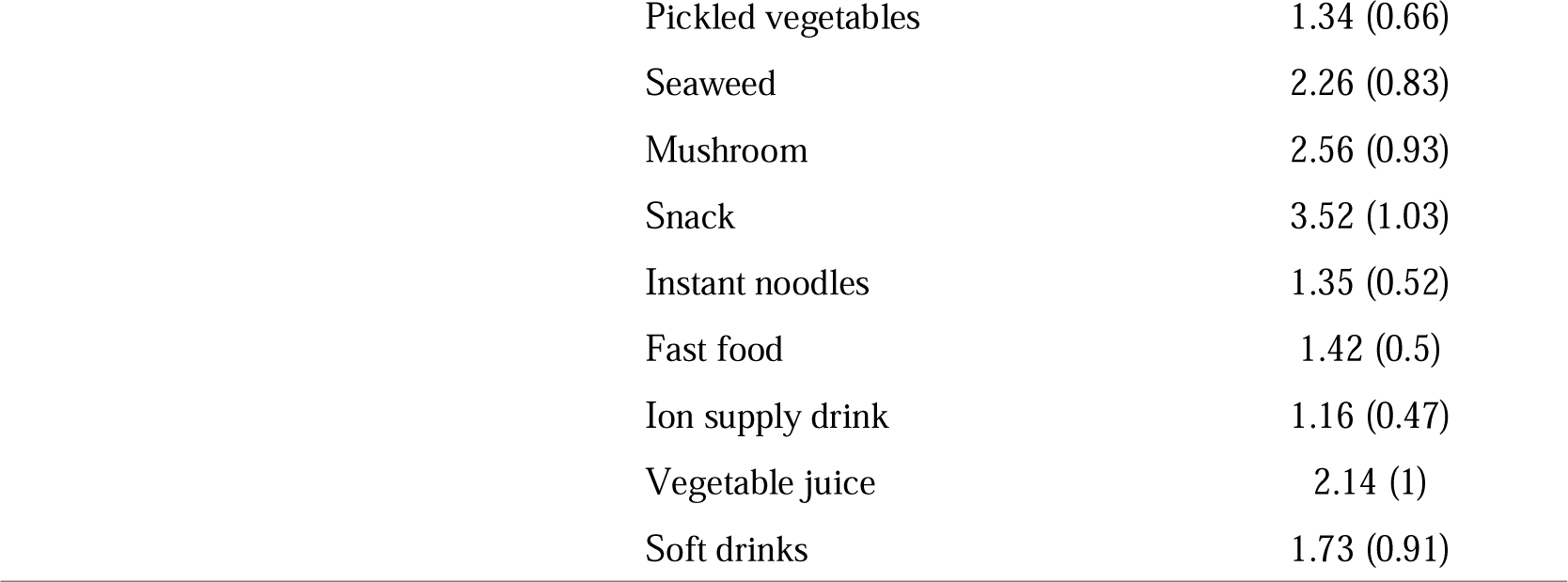
Characteristics of participants.

| Variable |  | N (%) or Mean (SD) |
| --- | --- | --- |
| Child sex | Boys | 172 (60.56%) |
|  | Girls | 112 (39.44%) |
| Age of child (months) |  | 46.44 (6.27) |
| Family income | Less than 3 million yen<br>(about 26,000 USD) | 32 (11.27%) |
|  | 3-5 million yen<br>(about 26,000-43,360 USD) | 76 (26.76%) |
|  | 5-7 million yen<br>(about 43,360-60,700 USD) | 80 (28.17%) |
|  | More than 7 million yen<br>(about 60,700 USD) | 75 (26.41%) |
|  | No answer | 21 (7.39%) |
| Age of mother (years) |  | 35.36 (4.65) |
| Maternal education (years) |  | 14.62 (1.95) |
| Maternal psychiatric history | No | 278 (97.89%) |
|  | Yes (already treated) | 6 (2.11%) |
| Antimicrobial use within the last three months | No | 236 (83.10%) |
|  | Yes | 37 (13.03%) |
|  | Not sure | 11 (3.87%) |
| Birth mode | Vaginal delivery | 226 (79.58%) |
|  | C-section | 58 (20.42%) |
| Frequency of food intake per week<br>(1 = never eaten, 2 = 1-3 times, 3 = 4-6 times, 4 = 1 time every day, 5 = more than two times every day) | Rice or bread | 4.64 (0.84) |
|  | Unrefined grains | 1.37 (0.92) |
|  | Root vegetables | 3.56 (1.06) |
|  | Green and yellow vegetables | 3.19 (1.12) |
|  | Light colored vegetables | 3.18 (1.14) |
|  | Fruit | 3.34 (1.09) |
|  | Potatoes | 2.75 (0.9) |
|  | Meat | 3.52 (0.98) |
|  | Seafood | 2.71 (0.91) |
|  | Eggs | 2.99 (0.98) |
|  | Milk or cheese | 3.65 (1.15) |
|  | Yogurt or probiotic drinks | 2.93 (1.07) |
|  | Soy products | 2.76 (1.01) |
|  | Beans | 2.11 (0.85) |
|  | Natto | 2 (0.9) |

|  |  |
| --- | --- |
| Pickled vegetables | 1.34 (0.66) |
| Seaweed | 2.26 (0.83) |
| Mushroom | 2.56 (0.93) |
| Snack | 3.52 (1.03) |
| Instant noodles | 1.35 (0.52) |
| Fast food | 1.42 (0.5) |
| Ion supply drink | 1.16 (0.47) |
| Vegetable juice | 2.14 (1) |
| Soft drinks | 1.73 (0.91) |

### 2.2. Temperament assessment

Child temperament was assessed using maternal reports from the Japanese version of the Children’s Behavior Questionnaire Short Form (CBQ-SF) [24]. The CBQ is a reliable measure used to assess children’s temperament [25]. The CBQ-SF consists of 92 items. For each item, mothers rated their child’s behavior in various daily situations observed in the last two weeks on a 7-point scale (1 = not observed at all to 7 = always observed). The questionnaire consists of three main dimensions and 15 subscales: *Negative Affectivity* (including *Anger*, *Distress*, *Fear*, *Sadness*, and *Falling Reactivity*; *Falling Reactivity* is inverse subscale), *Surgency/Extraversion* (including *Activity Level*, *Approach*, *High-Intensity Pleasure*, *Impulsivity*, and *Shyness*; *Shyness* is inverse subscale), and *Effortful Control* (including *Attention Focusing*, *Inhibitory Control*, *Low-Intensity Pleasure*, *Perceptual Sensitivity*, and *Smiling and Laughter*). *Negative Affectivity* refers to the general tendency associated with the expression of negative emotions and stress response. *Surgency/Extraversion* refers to the general tendency associated with expressions of positive emotions, reward-seeking, and activity levels. *Effortful Control* refers to the general tendency associated with the voluntary management of attention, inhibitory control, and activating behaviors as required. For detailed definitions of each subscale, see Table 2 based on the previous study [14,26].

**Table 2.** Scale definitions of the Childhood Behavior Questionnaire (CBQ).

| Scale | Definition |
| --- | --- |
| *Negative Affectivity* | General dimension associated with expression of negative emotion and defensive response to potential threat |
| *Anger* | Negative affect related to confinement, interruption of ongoing tasks or goal blocking |
| *Distress* | Negative affect related to sensory qualities of stimulation, including intensity, rate or complexity of light, sound, and texture |
| *Falling Reactivity* * | Rate of recovery from peak distress, excitement, or general arousal |
| *Fear* | Negative affect related to anticipated pain, distress and/or threat |
| *Sadness* | Negative affect, tearfulness or lowered mood related to physical state, disappointment, loss, and/or response to other's suffering |
| *Surgency/Extraversion* | General dimension associated with expressions of positive emotions, reward seeking, and activity level |
| *Activity Level* | Gross motor activity, including rate and extent of movement |
| *Approach* | Excitement in the anticipation of pleasurable activities |
| *High-intensity Pleasure* | Pleasure or enjoyment related to high stimulus intensity, rate, complexity, novelty, and incongruity |
| *Impulsivity* | Speed of response initiation |
| *Shyness* * | Slow or inhibited approach and/or discomfort in social situations |
| *Effortful Control* | General dimension associated with voluntarily management of attention, inhibitory control, and activate behaviour as required |
| *Attention Focusing* | Ability to transfer attentional focus from one object or task to another |
| *Inhibitory Control* | Capacity to suppress inappropriate actions or responses |
| *Low-Intensity Pleasure* | Pleasure or enjoyment related to low stimulus intensity, rate, complexity, novelty, and incongruity |
| *Perceptual Sensitivity* | Detection of slight, low-intensity stimuli from the external environment |
| *Smiling and Laughter* | Positive affect in response to changes in stimulus intensity, rate, complexity, and incongruity |

\* denotes invert scale of main dimension.

### 2.3. Fecal sample collection and DNA extraction

Fecal samples were collected using Mykinso fecal collection kits containing guanidine thiocyanate solution (Cykinso, Tokyo, Japan), transported at ambient temperature, and stored at 4°C. DNA extraction from the fecal samples was performed using an automated DNA extraction machine (GENE PREP STAR PI-480, Kurabo Industries Ltd, Osaka, Japan) according to the manufacturer’s protocol.

### 2.4. 16S rRNA gene sequencing

The detailed methods are described in a previous article [27]. Briefly, amplicons of the V1V2 region were prepared using the forward primer (16S_27Fmod: TCG TCG GCA GCG TCA GAT GTG TAT AAG AGA CAG AGR GTT TGA TYM TGG CTC AG) and reverse primer (16S_338R: GTC TCG TGG GCT CGG AGA TGT GTA TAA GAG ACA GTG CTG CCT CCC GTA GGA GT). The libraries were sequenced in a 250-bp paired-end run using the MiSeq Reagent Kit v2 (Illumina; 500 cycles).

### 2.5. Taxonomy assignment based on 16S rRNA gene sequencing

The detailed processing methods are described in a previous article [28]. Briefly, data processing and assignment using the QIIME2 pipeline (version 2020.8) [29] were performed as follows: (1) joining paired-end reads, filtering, and denoizing with a divisive amplicon denoizing algorithm (DADA2) and (2) assigning taxonomic information to each amplicon sequence variant (ASV) using a naive Bayes classifier in the QIIME2 classifier.

### 2.6. Bioinformatics analysis

Beta diversity metrics were calculated to measure differences in gut microbiome composition between individuals. The study used unweighted and weighted Unifrac distances as beta diversity. Unweighted Unifrac distance indicates beta diversity incorporating the presence/absence of taxa. Weighted Unifrac distance refers to beta diversity incorporating the relative abundance of taxa. Beta diversity metrics were calculated with QIIME2’s ‘diversity’ plugin.

Alpha diversity metrics were calculated to measure gut microbiome diversity within individuals. The study measured alpha diversity at the ASV level using the Shannon index, Chao1, Observed species, and Faith’s phylogenetic diversity. The Shannon index is a measure of richness and evenness. Chao1 estimates the total number of ASVs that would be observed with infinite sampling. Observed species was simply the observed number of ASVs per sample. Faith’s phylogenetic diversity is a phylogenetic measure of taxon richness and is expressed as the number of tree units observed in the sample. Alpha diversity metrics were calculated with QIIME2’s ‘diversity’ plugin.

The relative abundance of genera was calculated as the number of sequencing reads of each taxon in a sample standardized by the total number of sequences generated for that sample. We left sequence counts that we could not classify to the taxonomic level of interest as unclassified counts of the lowest level possible. Finally, log-transformed values of the relative abundance were used in the analysis.

Data manipulation, analyses, and graphics were performed using R and RStudio (ver. 4.2.1 and 2022.02.3, respectively). The following R packages were used for analyses: qiime2R (ver. 0.99.6), microbiome R (ver. 1.18.0), and phyloseq (ver. 1.40.0).

### 2.7. Statistical analysis

The following analysis is based on a previous study [12]. As a prior analysis, principal coordinate analysis of beta diversity and principal component analysis of alpha diversity were performed to generate principal coordinate/component scores for use in the main analysis. Regarding beta diversity, principal coordinate analysis was performed on unweighted Unifrac distance and weighted Unifrac distance separately. Principal coordinate analysis was performed with QIIME2’s ‘diversity’ plugin. Regarding alpha diversity, principal component analysis was performed on four alpha diversity indices: Shannon index, Chao1, Observed species, and Faith’s phylogenetic diversity. See Note S1 for the result of the principal coordinate/component analysis.

To investigate the association between temperament and gut microbiota, the main analysis was conducted in two steps. In the first step, the correlation between temperament and principal coordinates/components of diversity metrics was tested using Pearson’s correlation coefficient to determine the variables for the regression analysis. The following variables were used: all temperament scales (three main dimensions and 15 subscales), four beta diversity principal coordinates (Weighted/Unweighted Unifrac PCo1 and Pco2), and two alpha diversity principal components (Alpha PC1 and PC2).

As a final step, the study performed multiple regression analyses only on beta diversity principal coordinates and/or alpha diversity principal components that showed a significant correlation with temperament in the first step analysis. Multiple linear regression models were built for temperament as response variables, beta diversity principal coordinates and/or alpha diversity principal components as explanatory variables, and the child’s sex and age as covariates (See Note S1 for detail of variable selection for multiple linear regression models). All variables except the child’s sex were z-standardized prior to the multiple regression analysis, and a standardized partial regression coefficient was reported. The study performed multiple regression analysis using the R function ‘pairwise_association_test_multiple_regression’ in the analysis code from a previous study [12] (https://github.com/argossy/gmia_public), partially modified for this study.

In addition, for beta diversity principal coordinates that were significantly associated with temperament, the study calculated Spearman’s rank correlation coefficient between the principal coordinate and the relative abundance of each genus to understand which bacteria contributed to the principal coordinate. Similarly, for alpha diversity principal components that were significantly associated with temperament, the study calculated Pearson’s rank correlation coefficient between the alpha principal component and four alpha diversity indices to understand which alpha diversity index contributed to the principal component.

For all analyses, Statistical significance levels were set at *p* < 0.05. Data manipulation, analyses, and graphics were performed using R and RStudio (ver. 4.2.1 and 2022.02.3, respectively) for principal coordinate/component analyses. Similarly, for other statistical analysis, data manipulation, analyses, and graphics were performed using R and RStudio (ver. 4.1.2 and 2022.07.1+554, respectively). The following R packages were used for visualizations: ggplot2 (ver. 3.4.2), ggbeeswarm (ver. 0.7.1), and ggpubr (ver. 0.6.0), for visualizations.

## 3. Results

The study conducted correlation analyses between temperament (i.e., three main dimensions and 15 subscales) and principal coordinates/components of microbiota diversity metrics (i.e., Weighted/Unweighted Unifrac PCo1 and PCo2, Alpha PC1, and PC2) (see Note S1 for details of the principal coordinate/component metrics). The results of the correlation analysis between temperament and beta diversity principal coordinates showed that Unweighted Unifrac principal component 2 (Unweighted Unifrac PCo2) had negative correlations with *Negative Affectivity* (*r* = -0.17, *p* = 0.003, Figure 1A), and with its subscales *Anger* (*r* = -0.12, *p* = 0.035, Figure 1B), *Fear* (*r* = -0.12, *p* = 0.044, Figure 1C), and *Sadness* (*r* = -0.22, *p* < 0.001, Figure 1D). *Shyness*, an inverted subscale of *Surgency/Extraversion*, was also negatively correlated with Unweighted Unifrac PCo2 (*r* = -0.16, *p* = 0.009, Figure 1E). Conversely, Unweighted Unifrac Pco2 had positive correlations with *Surgency/Extraversion* (*r* = 0.14, *p* = 0.019, Figure 2A) and its subscale *Impulsivity* (*r* = 0.12, *p* = 0.043, Figure 2B). These results persisted even after controlling for covariates (i.e., sex and age) in the multiple regression analysis (*Negative Affectivity β* = -0.17, *p* = 0.004; *Anger β* = -0.12, *p* = 0.045; *Fear β* = -0.12, *p* = 0.0498; *Sadness β* = -0.22, *p* < 0.001 and *Shyness β* = -0.16, *p* = 0.008, Figure 1f; *Surgency/Extraversion β* =0.15, *p* = 0.013; *Impulsivity β* =0.13, *p* = 0.025, Figure 2D).

**Figure 1.**
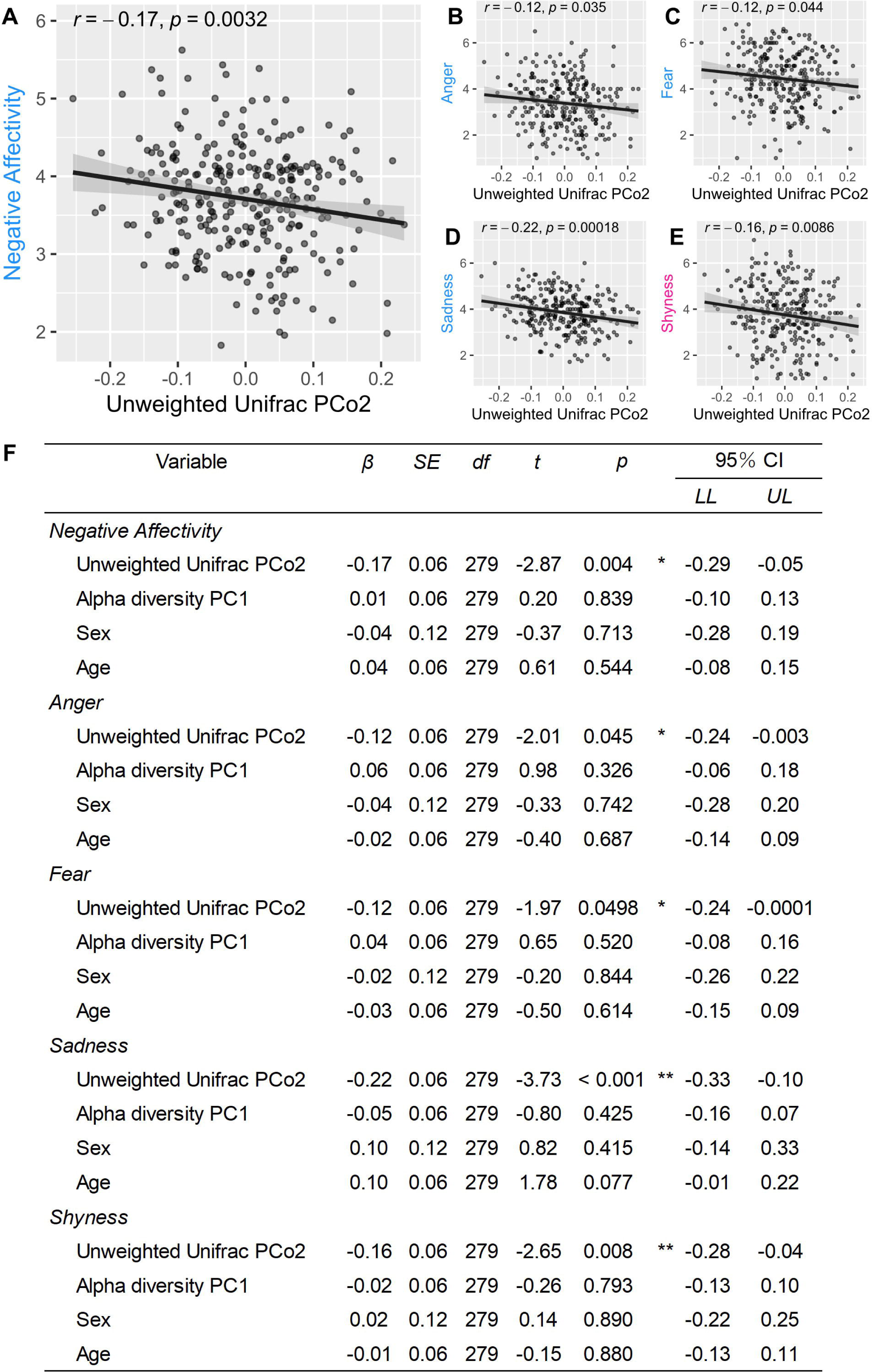
The relationships between the gut microbiome and *Negative Affectivity*, *Anger, Fear*, *Sadness*, and *Shyness*. (A) to (E) present scatterplots illustrating the associations between Unweighted Unifrac PCo2 and each temperament dimension. *Anger*, *Fear*, and *Sadness* are subscales of *Negative Affectivity*, while *Shyness* is the inverted subscale of *Surgency/Extraversion*. (F) shows the statistical results of multiple regression analysis. The abbreviation ‘CI’ refers to confidence interval, ‘*LL*’ represents the lower limit, and ‘*UL*’ denotes the upper limit. * *p* <.05 and ** *p* <.01.

**Figure 2.**
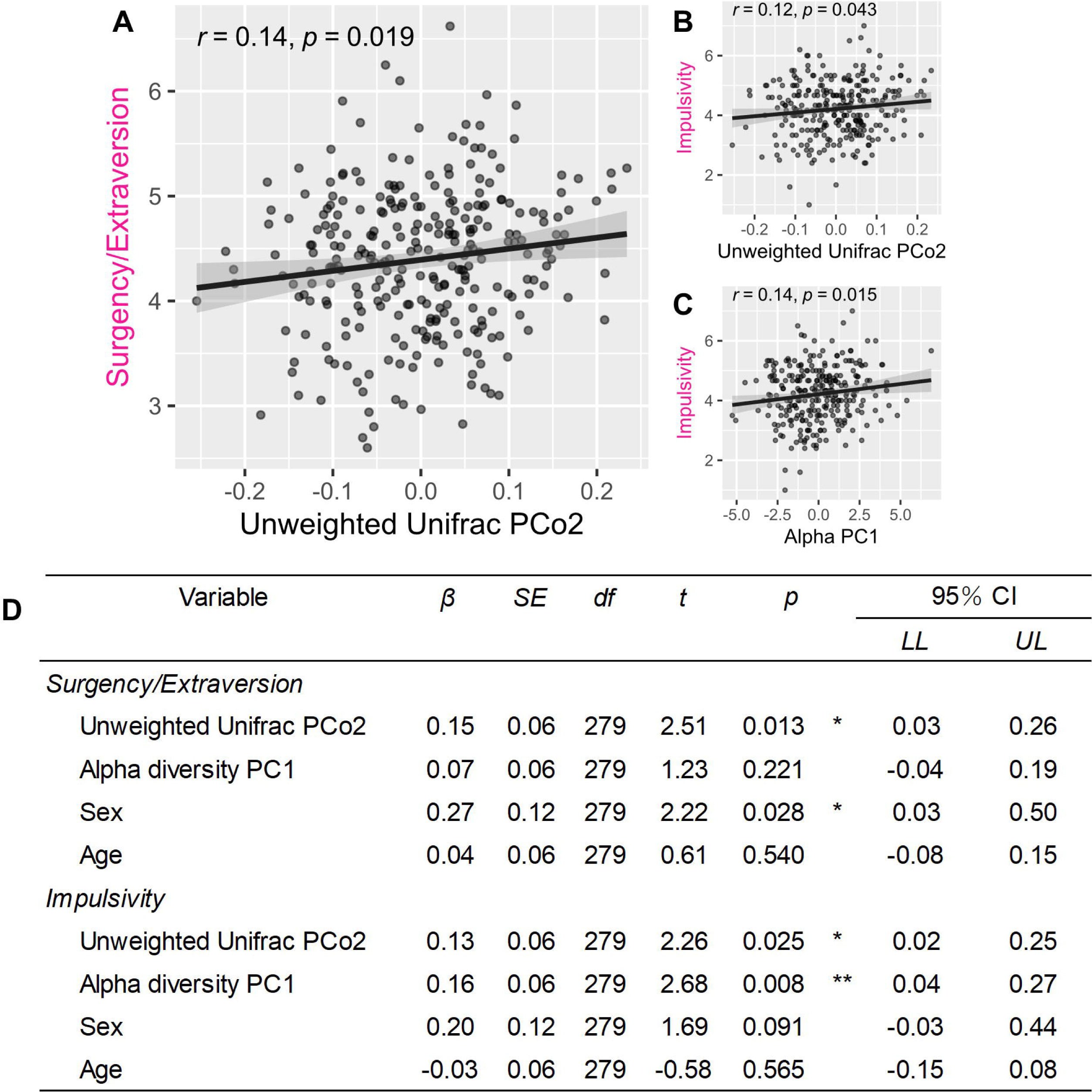
The relationships between gut microbiome and *Surgency/Extraversion* and *Impulsivity*. (A) and (B) show the associations between Unweighted Unifrac PCo2 and *Surgency/Extraversion* and *Impulsivity*, respectively. (C) shows the association between Alpha PC1 and *Impulsivity*. (D) shows the statistical results of multiple regression analysis. The abbreviation ‘CI’ represents the confidence interval, ‘*LL*’ refers to the lower limit, and ‘*UL*’ denotes the upper limit. * *p* <.05 and ** *p* <.01.

To gain a deeper understanding of these results, the study analyzed which bacteria made significant contributions to Unweighted Unifrac PCo2. Unweighted Unifrac PCo2 was positively correlated with the relative abundance of *Haemophilus* (*rs* = 0.51, *p* < 0.001), *Faecalibacterium* (*rs* = 0.33, *p* < 0.001), *Erysipelotrichaceae* (*rs* = 0.31, *p* < 0.001), and *Collinsella* (*rs* = 0.31, *p* < 0.001); and negatively correlated with the relative abundance of *Eisenbergiella* (*rs* = -0.63, *p* < 0.001), *Coprobacillus* (*rs* = -0.61, *p* < 0.001), *Flavonifractor* (*rs* = -0.59, *p* < 0.001), *UBA1819* (*rs* = -0.58, *p* < 0.001), *Anaerotruncus* (*rs* = -0.55, *p* < 0.001), *Tuzzerella* (*rs* = -0.53, *p* < 0.001), and *Eggerthella* (*rs* = -0.52, *p* < 0.001) (Figure S1). These findings indicate that higher values of Unweighted Unifrac PCo2 are associated with an increased abundance of *Haemophilus*, *Faecalibacterium*, *Erysipelotrichaceae*, and *Collinsella*; and lower abundance of *Eisenbergiella*, *Coprobacillus*, *Flavonifractor*, *UBA1819*, *Anaerotruncus*, *Tuzzerella*, and *Eggerthella*.

The results of correlation analysis between temperament and alpha diversity principal components showed that Alpha diversity principal component 1 (Alpha PC1) had a positive correlation with *Impulsivity* (*r* = 0.14, *p* = 0.015, Figure 2C). This persisted after controlling for covariates in the multiple regression analysis (*β* = 0.16, *p* = 0.008, Figure 2D). The higher values of Alpha PC1 indicate greater bacterial and taxonomic richness as well as bacterial evenness (Figure S2). Therefore, the results suggest that children who have higher bacterial and taxonomical abundance, as well as evenness have higher *Impulsivity* temperament.

See Note S1 for details on variable selection for the multiple linear regression models. See Figure S3 for all results, including null results for correlation analysis between temperament, beta diversity principal components, and alpha diversity principal components.

## 4. Discussion

The study investigated the association between temperament and gut microbiota in preschool children. Most importantly, significant associations between the composition of the gut microbiota, as assessed by Unweighted Unifrac PCo2, and *Negative Affectivity*, *Surgency/Extraversion*, as well as specific subscales within these dimensions were established. Negative association was found between Unweighted Unifrac PCo2 and *Negative Affectivity*, as well as its subscales *Fear*, *Anger*, and *Sadness*. Furthermore, the study found a negative association between Unweighted Unifrac PCo2 and *Shyness*, which represents the inverse subscale of *Surgency/Extraversion*. In contrast, Unweighted Unifrac PCo2 exhibited a positive association with *Surgency/Extraversion* and its subscale, *Impulsivity*.

*Negative Affectivity* reflects the general tendency to express negative emotions and encompasses individual differences in stress responses. *Surgency/Extraversion*, by contrast, reflects reactivity to potentially rewarding stimuli and environments. This dimension encompasses traits such as high positive emotional expression and increased reward-seeking (for detailed definitions of each subscale, see Table 2). The results therefore suggest that children with higher Unweighted Unifrac PCo2 are less likely to display negative emotions and stress reactions. Instead, they are more likely to exhibit positive emotions and actively engage with their surroundings, including interacting with others and demonstrating interest in novel stimuli.

Considering the functional aspects of bacteria associated with Unweighted Unifrac PCo2, it is probable that this value is linked to inflammation/anti-inflammation processes. Specifically, the bacteria that showed a positive correlation with Unweighted Unifrac PCo2 included *Faecalibacterium*, a butyrate-producing bacterium known for its anti-inflammatory effect [30,31]. However, bacteria exhibiting a negative correlation with Unweighted Unifrac PCo2 included *Eggerthella*, which is associated with gastrointestinal inflammation [32], and *Flavonifractor*, which potentially inhibits antioxidant and anti-inflammatory effects [33,34]. Therefore, a higher Unweighted Unifrac PCo2 value may indicate a greater presence of anti-inflammatory bacteria and a lower abundance of inflammatory bacteria. In summary, children with more anti-inflammatory bacteria and lower levels of inflammatory bacteria (higher Unweighted Unifrac PCo2) are less likely to exhibit negative emotions and stress reactions (lower *Negative Affectivity*), while they are more likely to display positive emotions and engage with their environment (higher *Surgency/Extraversion*) (Figure 3).

**Figure 3.**
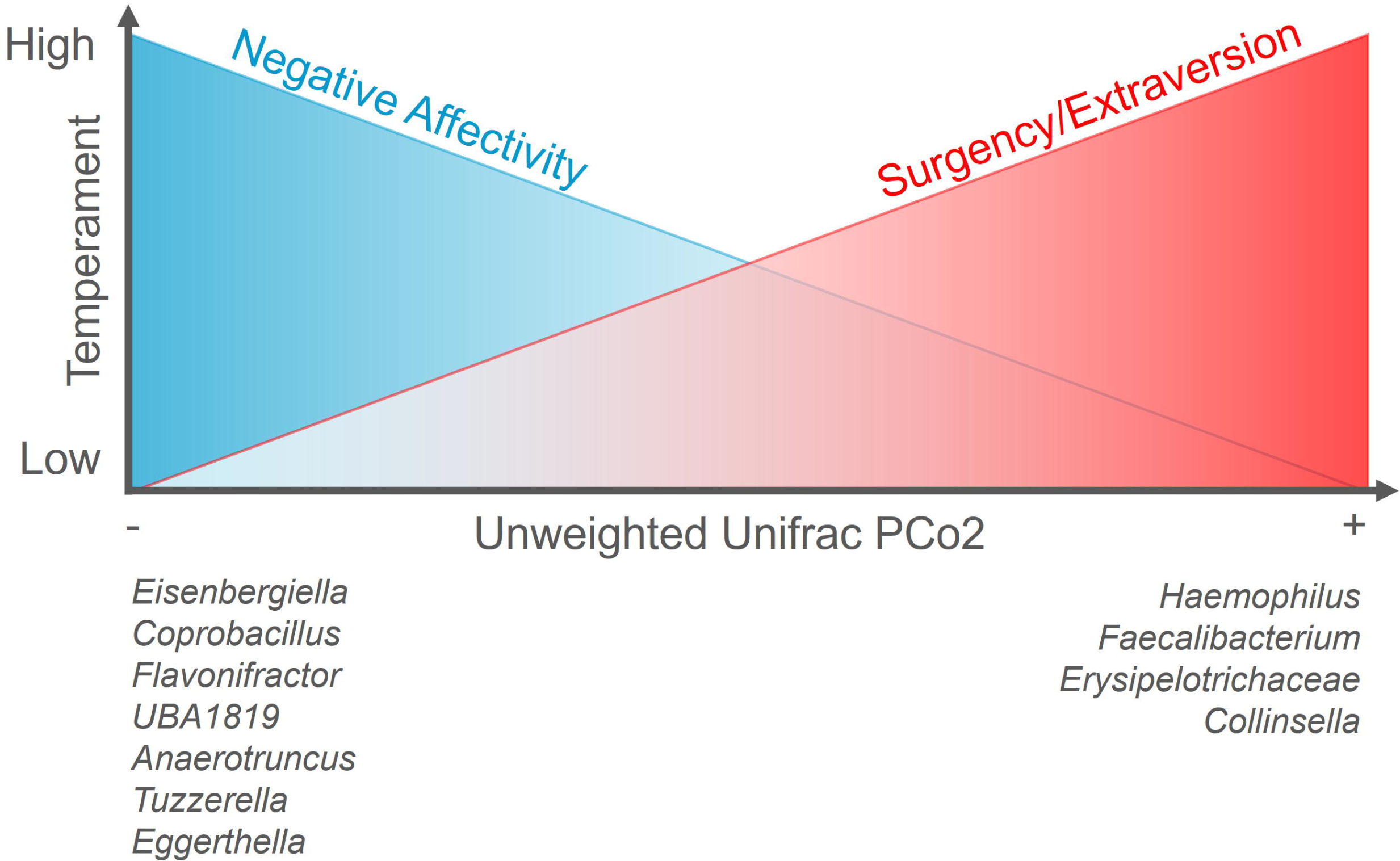
Conceptual image depicting the association between *Negative Affectivity*, *Surgency/Extraversion*, and gut microbiota composition. A higher score on Unweighted Unifrac PCo2 (indicating a higher abundance of anti-inflammatory bacteria and fewer inflammatory bacteria) is associated with higher levels of *Surgency/Extraversion* and lower levels of *Negative Affectivity*. Conversely, a lower score on Unweighted Unifrac PCo2 (indicating a higher abundance of inflammatory bacteria and fewer anti-inflammatory bacteria) is associated with higher levels of *Negative Affectivity* and lower levels of *Surgency/Extraversion*.

Additionally, the study found that the diversity of the gut microbiota, as measured by Alpha PC1, showed a positive association with *Impulsivity*, a subscale of *Surgency/Extraversion*. *Impulsivity* reflects the tendency to actively approach and interact with the environment and novel stimuli. Therefore, this result suggests that children with higher Alpha PC1 are more actively engaged in interacting with their surroundings. It is important to note that higher values of Alpha PC1 indicate greater values for all four indices of alpha diversity, which means a higher bacterial richness, taxonomic richness, and bacterial evenness within the gut microbiota. In general, alpha diversity is considered an indicator of gut microbiota maturity, as it tends to increase with age in children under 5 years of age [4]. Thus, the results suggest that children with a more mature gut microbiota are more likely to actively engage with their environment.

The findings of this study, particularly the relationship observed between gut microbiota and temperament, as depicted in Figure 3, have important implications for predicting the developmental outcomes of children’s mental health. Children positioned at the left end of Figure 3 demonstrate a temperament characterized by high levels of *Negative Affectivity* and low levels of *Surgency/Extraversion*. Children with these temperament traits typically exhibit greater negative emotional expression and stress reactions while displaying less positive emotional expression and social interaction. These specific temperamental characteristics are associated with an increased likelihood of experiencing internalizing symptoms such as anxiety and depression during late childhood and adolescence [16–18]. In other words, children located on the left side of Figure 3 have temperamental traits that put them at a higher risk of developing symptoms such as anxiety and depression.

Furthermore, concerning the gut microbiota, children positioned at the left end of Figure 3 exhibit higher levels of inflammatory bacteria and lower levels of anti-inflammatory bacteria. Inflammatory bacteria, such as *Eggethella*, have been associated with psychiatric disorders like anxiety [35] and depression [36]. Conversely, bacteria with anti-inflammatory properties, like *Faecalibacterium*, show potential as probiotics for psychiatric disorders [37]. A recent review study reported that depressed patients had elevated levels of *Eggerthella* and reduced levels of *Faecalibacterium* compared to healthy controls [6,38]. That is, children positioned on the left side of Figure 3 have a higher abundance of inflammatory bacteria, which are linked to mental problems such as anxiety and depression while having a lower presence of anti-inflammatory bacteria, which could potentially have a positive impact on mental health.

To summarize, children depicted on the left side of Figure 3 may have characteristics associated with a heightened risk of psychiatric disorders, considering both temperament and gut microbiota factors. Therefore, it is vital to closely monitor the developmental progress of children displaying these specific temperament and gut microbiota characteristics. Additionally, it is crucial to explore potential microbial intervention strategies based on the assessment of children’s gut microbiota patterns, particularly during periods when the microbiota is highly malleable. Dietary interventions, such as incorporating probiotics, have the potential to mitigate risks and support positive developmental trajectories. The findings underscore the significance of considering the relationship between gut microbiota and temperament in the context of mental health outcomes. They emphasize the necessity of early assessment and intervention strategies targeting the gut microbiota to potentially mitigate the risk of mental health disorders and promote favorable developmental outcomes in children. Subsequently, the study discusse the consistency and differences between the results of our study in preschool children and those of previous studies conducted on infants and toddlers. First, it compared the bacteria associated with *Surgency/Extraversion* observed in this study with those identified in previous studies. The results did not align with prior findings for infants [19,39] and toddlers [40]. However, considering the general developmental transition of gut microbiota, it is plausible that *Surgency/Extraversion* may be associated with a higher abundance of bacteria that are commonly observed during specific developmental periods. For instance, a study demonstrated that the relative abundance of *Bifidobacterium* in the neonatal period (1-3 weeks after birth) was positively associated with *Surgency/Extraversion* at one year of age [19]. Similarly, another study found that the abundance of *Streptococcus* at 2.5 months was positively associated with *Surgency/Extraversion* at 6 months [39]. *Streptococcus* and *Bifidobacterium* are lactic acid bacteria whose abundance tends to peak before the age of one year and subsequently declines [4]. In the current study, *Faecalibacterium* showed positive associations with *Surgency/Extraversion* and its subscales. *Faecalibacterium* is a type of bacteria that increases after the age of one year and stabilizes around 3–5 years of age [4]. Therefore, having bacteria that are generally more abundant during specific periods, reflecting a well-developed bacterial flora, may be associated with *Surgency/Extraversion*.

Second, the focus shifted toward the association between *Surgency/Extraversion* and the diversity of gut microbiota. The study found a positive association between alpha diversity and *Impulsivity*, a subscale of *Surgency/Extraversion*, which aligns with a previous study conducted on toddlers [40]. However, this finding differs from studies conducted on infants during their first year of life [19, 39], where no significant association between alpha diversity and *Surgency/Extraversion* was observed. It suggests that the relationship between *Surgency/Extraversion* and microbiome diversity may become clearer after the children reach 1.5 years of age. In summary, the study’s findings regarding the association between gut microbiota composition/diversity and *Surgency/Extraversion* in preschool children differs from those of previous infant studies. These differences may be attributed to the developmental transitions in gut microbiota and the specific time periods when certain bacterial abundances peak. Further research is required to investigate this association across different age groups and gain a more comprehensive understanding of the intricate dynamics between gut microbiota, temperament, and early childhood development.

Furthermore, the study compared the findings from our study with those of previous studies in terms of the association between *Negative Affectivity* and gut microbiota. The bacteria associated with *Negative Affectivity* in our study were not consistent with previous research on infants [12,19,39,41] and toddlers [40]. Moreover, there is inconsistency among previous studies in identifying the bacteria associated with *Negative Affectivity*. For example, an infant study showed that lower levels of *Bacteroides* and higher levels of *Lactobacillus* and *Bifidobacterium* were associated with increased fearful behavior toward novel non-social stimuli at one year of age [12]. However, another study with the same age group found that a higher abundance of *Lactobacillus* was associated with lower levels of *Negative Affectivity* [19]. *Lactobacillus* and *Bifidobacterium* are both types of lactic acid bacteria. There was no significant association between lactic acid bacteria and *Negative Affectivity* in the toddler study [40] or the current study. Therefore, while some association between lactic acid bacteria and *Negative Affectivity* is observed during infancy, it may change during early childhood. Furthermore, the association between *Negative Affectivity* and gut microbiota diversity was compared between our study and previous studies. The study found no significant association between alpha diversity and *Negative Affectivity*, but some infant studies have reported an association between alpha diversity and *Negative Affectivity* or its subscale *Fear* [12,39]. Thus, the association between gut microbiome diversity and *Negative Affectivity* may be observed only in infancy.

The results of the study have two implications for further research. First, the association between temperament and gut microbiota observed (i.e., lower levels of anti-inflammatory bacteria and higher levels of inflammatory bacteria associated with low *Surgency/Extraversion* and high *Negative Affectivity*) overlaps with important factors that predict mental health risk. Longitudinal studies are required to investigate whether these characteristics at age 3–4 affect developmental outcomes at the age of 5–6, which is a critical period in the development of executive functions involved in emotion regulation and the risk of mental problems. In this regard, another of our studies [42] has already found that children at risk for emotional control have higher levels of inflammation-associated bacteria. Second, comparing the current study to previous research involving infants reveals the importance of exploring the relationship between gut microbiota and temperament while considering alterations in gut microbiota composition during infancy and childhood. However, the majority of studies have primarily focused on examining the association between gut microbiota and temperament in American-European infants. Hence, cross-sectional studies are essential to determine whether the associations between temperament and gut microbiota, as demonstrated in previous infant studies, can be replicated in infants from diverse countries. Additionally, it remains unclear how gut microbiota in infancy is related to both gut microbiota and temperament at the age of 3–4. To address these questions, we have already initiated a longitudinal cohort study of a Japanese population.

## 5. Conclusions

This study revealed that gut microbiota composition related to gut inflammation/anti-inflammation was associated with positive/negative emotional expression, proximity to rewards, and behavioral inhibition tendencies. Furthermore, gut microbiota maturity was associated with higher levels of active approach and exploration. Although this study does not prove causality, it contributes to deepening the understanding of the biological mechanisms behind psychological traits in humans. It is also important to note that the temperamental traits associated with gut microbiota in this study overlap with important factors that predict mental health risk. Designing potential microbial intervention strategies such as specific probiotic supplementation is also expected to prevent and ameliorate psychological/emotional problems from an early developmental stage in children.

## Acknowledgments

This research was funded by Grant-in-Aid for Scientific Research from Japan Society for the Promotion of Science (JPSP), grant numbers 17H01016, 19K21813, and 21H04981 to M.M. (Masako Myowa); the Japan Science and Technology Agency (JST) Moonshot Research and Development Program, grant number JPMJMS2296 to M. M. (Masako Myowa); JST Center of Innovation Program, grant number JPMJCE1307 to M. M. (Masako Myowa); Kieikai Research Foundation to M. M. (Masako Myowa); Grant-in-Aid for JSPS Fellows, grant numbers 22KJ1868 to E.U., 19J15173 and 22J01448 to M.M. (Michiko Matsunaga).

