## Supplementary Material for "Temperament in early childhood is associated with gut microbiota composition and diversity"

### Note S1. Results of principal coordinate/component analysis and variable selection for multiple linear regression models.

#### Results of beta diversity principal coordinate analysis

The results of the principal coordinate analysis performed on weighted Unifrac distance showed that principal coordinate 1 (Weighted Unifrac PCo1) explained 13% of the variance, while principal coordinate 2 (Weighted Unifrac PCo2) explained 8% of the variance. Similarly, the results of the principal coordinate analysis performed on the unweighted Unifrac distance indicated that principal coordinate 1 (Unweighted Unifrac PCo1) explained 13% of the variance, while principal coordinate 2 (Unweighted Unifrac PCo2) explained 5% of the variance. For both weighted and unweighted Unifrac distance, the inclusion of principal coordinate 3 did not change the cumulative contribution much. Therefore, we used up to PCo2 for subsequent analyses.

#### Results of alpha diversity principal component analysis

Principal component analysis performed on four alpha diversity indices showed that principal component 1 (Alpha PC1) explained 92% of the variance, while principal component 2 (Alpha PC2) explained 6% of the variance. One reason for the high variance explanation rate of Alpha PC1 is that strong positive correlations were observed between Alpha PC1 and all four alpha diversity indices (Supplementary Figure 2). Up to PC2 for subsequent analyses were used.

#### Variable selection for multiple linear regression models

The study used correlation analysis between temperament and principal coordinates/components of diversity metrics to determine the variables of the multiple regression analysis. Only principal coordinates/components that showed significant correlations with temperament in this analysis were used in regression analysis. Unweighted Unifrac PCo2 was negatively correlated with *Negative Affectivity* ($r$ = -0.17, $p$ = 0.003), *Anger* ($r$ = -0.12, $p$ = 0.035), *Fear* ($r$ = -0.12, $p$ = 0.044), *Sadness* ($r$ = -0.22, $p$ < 0.001), and *Shyness* ($r$ = -0.16, $p$ = 0.009), and positively correlated with *Surgency/Extraversion* ($r$ = 0.14, $p$ = 0.019) and *Impulsivity* ($r$ = 0.12, $p$ = 0.043). Alpha PC1 was positively correlated with *Impulsivity* ($r$ = 0.14, $p$ = 0.015). Thus, Unweigted Unifrac PCo2 and Alpha PC1 were selected as explanatory variables in regression analysis. Unweighted Unifrac PCo1 was positively correlated with *Impulsivity* ($r$ = 0.117, $p$ = 0.0495). However, since Unweighted Unifrac PCo1 also had a strong positive correlation with Alpha PC1 ($r$ = 0.9, $p$ < 0.001), Unweighted Unifrac PCo1 was excluded from the explanatory variables in the regression model due to multicollinearity. No significant correlation existed between temperament and Weighted Unifrac PCo1, Weighted Unifrac PCo2 as well as Alpha PC2, so we excluded these variables from the regression analyses. See Supplementary Figure 3 for all results, including null results for correlation analysis.

Based on previous research, the study included the child's sex and age as covariates in the regression model, considering their potential influence on both temperament and gut microbiota. Sex differences have been observed in both temperament [1] and gut microbiome [2]. Indeed, previous studies investigating the relationship between temperament and gut microbiota in infants [3] and toddlers [4] have suggested the importance of sex as a covariate. Regarding age, temperament is considered to be a relatively stable trait, but it does undergo some changes during development [5]. Gut microbiome composition and diversity also change drastically up to around the age of 5 [6]. Therefore, child age was considered an important covariate in this study.

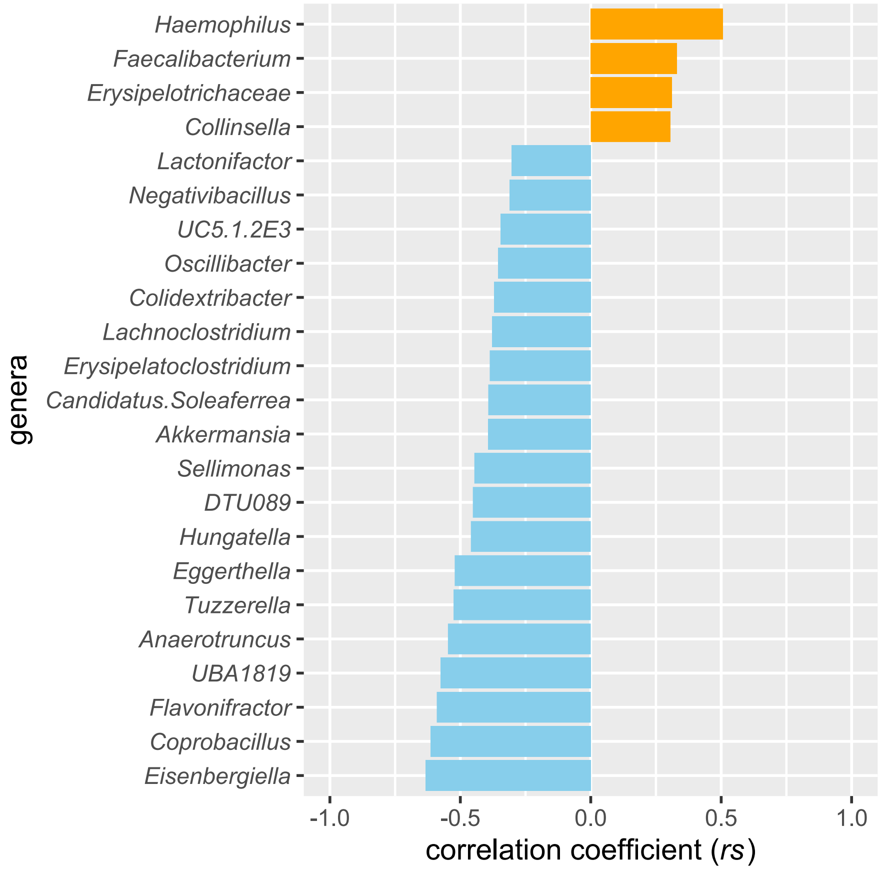


**Figure S1. Correlation coefficient between Unweighted Unifrac PCo2 and relative abundance of each genus.** Genera with medium to large effect sizes (i.e. correlation coefficients above 0.03) were selected for inclusion in this figure.

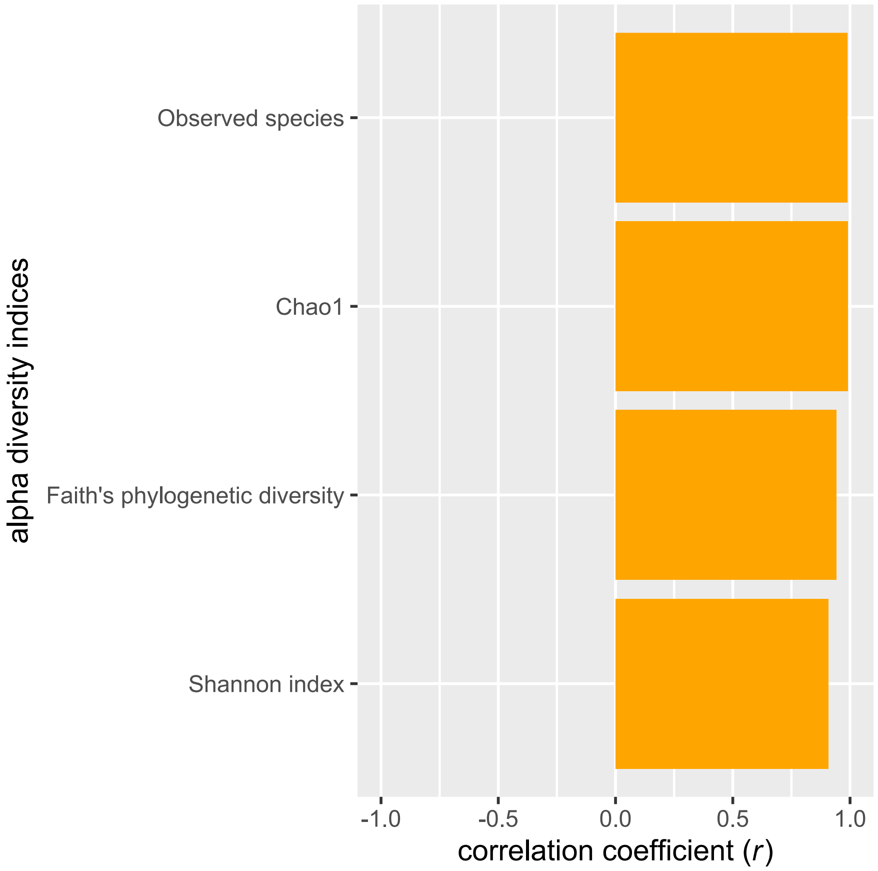


**Figure S2. Correlation coefficient between Alpha PC1 and each alpha index.**

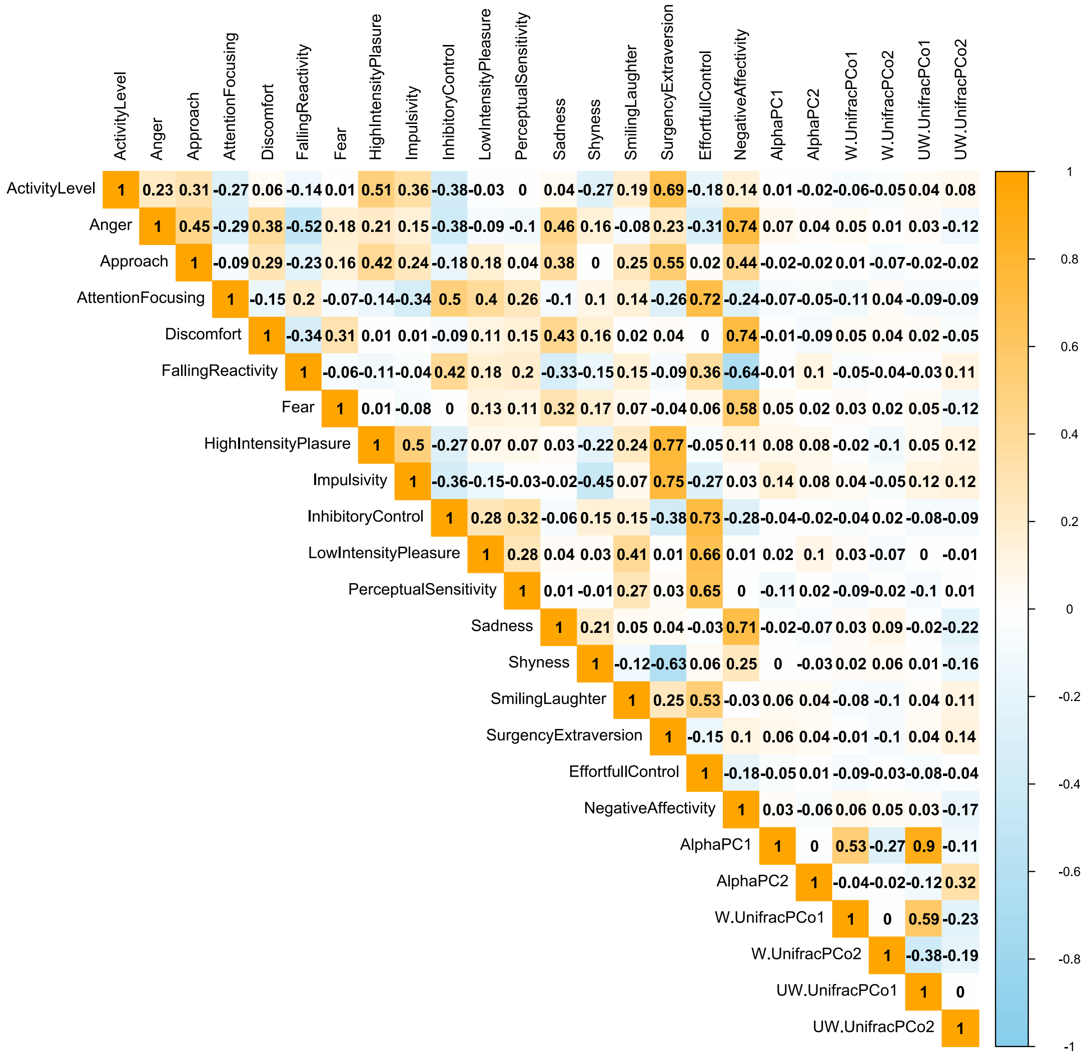


**Figure S3. Correlation coefficients between temperament, beta diversity principal coordinates, and alpha diversity principal components.** Temperament scales: three main dimensions, 15 subscales; beta diversity principal coordinates: Weighted/Unweighted Unifrac PCo1 and PCo2; alpha diversity principal components: Alpha PC1 and PC2.
